# Glucocorticoid signaling delays castration-induced regression in murine models of prostate cancer

**DOI:** 10.1101/2021.10.11.463722

**Authors:** Aerken Maolake, Renyuan Zhang, Kai Sha, Shalini Singh, Chunliu Pan, Bo Xu, Gurkamal Chatta, Michalis Mastri, Kevin H. Eng, John J. Krolewski, Kent L. Nastiuk

## Abstract

Androgen deprivation therapy (ADT) induces regression of recurrent and advanced prostate cancer (PrCa), but many tumors recur. To understand the response to ADT, changes in tumor volume were imaged after castration of murine PrCa models. While mouse (non-tumor) prostate begins to regress within two days of castration, murine PrCa regresses after a delay of 3-14 days in two distinct mouse models. Intra-tumoral androgens are undetectable after castration, but tumor cells proliferate during this period. Intratumoral glucocorticoids and glucocorticoid receptor (GR) protein increase, as does GR mRNA and a set of GR-regulated genes specifically in tumor epithelial cells identified using scRNAseq. A selective GR antagonist (CORT125281, relacorilant), in clinical trials for late-state PrCa, eliminates the delayed regression phenotype in both models. Thus, activated GR signaling and murine tumor proliferation following castration resembles the GR-dependent escape mechanism of castrate resistant PrCa. These results suggest simultaneous inhibition of GR and androgen receptor signaling could improve PrCa therapy.

**In brief:** Androgen deprivation therapy for high risk and recurrent prostate cancers is initially effective, but ultimately fails; better understanding the mechanisms should improve therapy. In two murine prostate cancer models, GR signaling is activated immediately following castration, substituting for the acute reduction in AR signaling, and allowing for continued tumor growth. This continued growth is blocked by relacorilant, selective GR antagonist in clinical trials for late-state PrCa.

**Highlights:**

- Androgen deprivation therapy induces regression of prostate cancer, but tumors recur
- Murine PrCa continues to proliferate for 3-14 days in two distinct mouse prostate cancer models
- Tumor cells proliferate during this period, and intratumoral glucocorticoids and glucocorticoid receptor (GR) protein increase, as does GR mRNA and a set of GR-regulated genes
- Relacorilant, a selective GR antagonist in clinical trials for late-state PrCa, eliminates the delayed regression

## INTRODUCTION

Prostate cancer (PC) is one of the more frequently diagnosed malignancies in U.S. men with 191,930 new diagnoses and 33,330 deaths in 2020 (Siegel et al., 2020). While many tumors are indolent or can be eliminated by curative surgery or radiotherapy, PC that presents at an advanced stage, or that is recurrent or metastatic is difficult to treat and cure. These tumors eventually constitute the bulk of prostate cancer deaths (Teo et al., 2019). Although the precise etiology of PC remains uncertain, primary tumor growth is driven by the androgen receptor (AR) axis, via re-programming of the AR cistrome (Pomerantz et al., 2015). The re-programmed AR cistrome is, in turn, dependent on the presence of circulating androgens, providing the rationale for androgen deprivation therapy (ADT). ADT has long been the mainstay therapy for PC. It is initially effective but most prostate cancers become resistant within a few years and PC progresses to castration-resistant prostate cancer (CRPC). Attempts to improve PC therapy by combining ADT with either chemotherapy, androgen biosynthesis inhibitors or AR-targeting anti-androgens are ongoing (Sartor and de Bono, 2018). Thus, improved understanding of the mechanism of tumor response to ADT may be valuable in interpreting the results of clinical trials.

To investigate the response of prostate cancers to ADT, we have employed two well-characterized genetically engineered murine models (GEMMs) of prostate cancer, driven by prostate-specific PTEN-loss or c-MYC gain, respectively (reviewed in ref. (Arriaga and Abate-Shen, 2019)). These genetic lesions are common in both primary and metastatic human prostate cancers, suggesting the corresponding mice are robust models of human prostate cancer. To optimize our understanding of tumor development and response to therapy, we have developed a protocol to serially image murine prostates using high frequency ultrasound (HFUS) in combination with software that enables three-dimensional reconstruction from the ultrasound image data (Singh et al., 2015). This relatively rapid and affordable methodology can accurately determine tumor size and location within the multi-lobed murine prostate and non-invasive serial imaging provides insight into the kinetics of tumor volume under various conditions or treatments.

In this report, we demonstrate that castration causes the tumors in these murine models of prostate cancer to regress over a period of 4 weeks, but that there is an initial delay in regression, relative to the effect of castration on the normal murine prostate (Davis et al., 2011). We combined molecular analyses with the use of a selective glucocorticoid receptor (GR) antagonist (Kach et al., 2017) to demonstrate that delayed regression occurs because GR signaling transiently substitutes for AR signaling in the prostate cancers.

## RESULTS

### Ultrasound imaging precisely measures prostate tumor volume changes in GEMMs

We have previously established the precision and reproducibility of magnetic resonance (MR) (Nastiuk et al., 2007) and high frequency ultra-sound (HFUS) (Singh et al., 2015) imaging protocols for monitoring the volume of normal prostate as well as prostate cancers (Pan et al., 2020) in murine models. We compared the utility of HFUS and MR imaging by sequentially imaging three mice bearing MYC driven prostate cancers with both modalities. Both HFUS and MR imaging showed a high degree of correlation for tumor volume determination (Supplementary Fig. S1). MR imaging has a higher resolution, and it is therefore easier to visually distinguish tumor boundaries, but image capture itself is slower and more expensive. For HFUS imaging, image capture is faster and the three-dimensional nature of the HFUS data in combination with AMIRA software allows semi-automated segmentation to determine tumor volume (6), facilitating serial measurements. Therefore, we employed the HFUS methodology in these studies to maximize data acquisition (number of mice and number of volume determinations per mouse).

### Castration induces delayed regression in a PTEN-deficient mouse model

To determine the effects of castration on primary prostate cancers, we surgically castrated groups of mice bearing PTEN-deficient prostate cancers, and serially imaged each mouse using HFUS to determine the change in tumor volume over time. In the male mice of this GEMM, loss of PTEN expression occurs - at puberty - in the prostate epithelial cells and leads to the development of PTEN-deficient prostate cancers, with essentially 100% penetrance (Wang et al., 2003). Fig. 1 demonstrates that, following castration, PTEN-deficient tumors continue to increase in volume for seven days, before beginning to regress. During the initial 7 days post-castration, the average volume of the tumors in the castrated mice were indistinguishable from sham-operated mice (Fig. 1B), suggesting that the prostate tumors were growing at the same rate in castrated and intact hosts. We tracked the tumor volume in multiple cohorts of the same GEMM, that differed in average initial tumor volume (Fig. 2A-D). Onset of castration-induced regression was delayed in 51/55 tumors, and the length of the delay correlated with the initial volume of the tumor (Fig. 2E).

**Figure 1.**
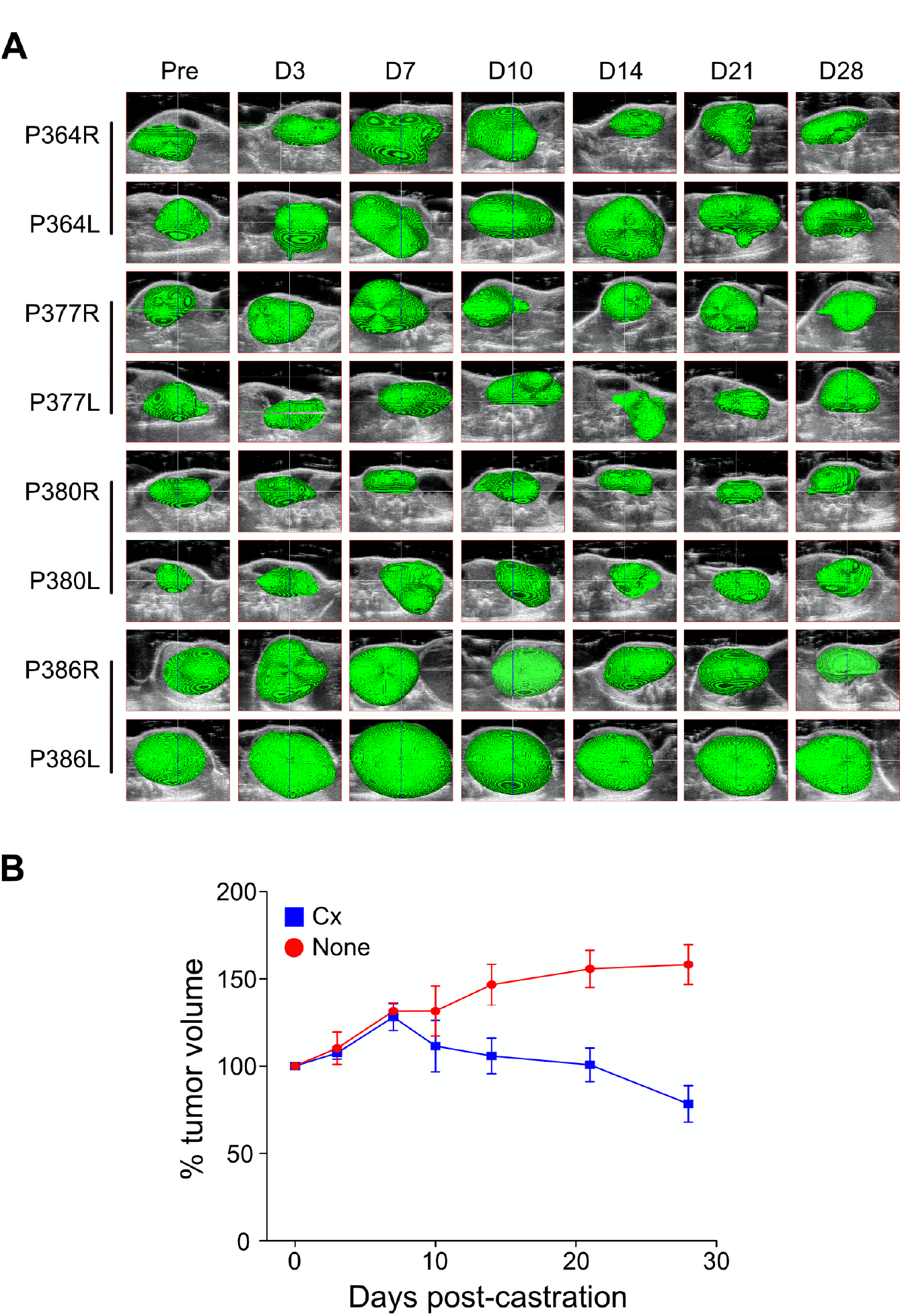
Castration induces delayed regression of PTEN-deficient prostate cancers. Tumor-bearing PB-Cre4:Pten^fl/fl^ mice were sham-operated (None) or castrated (Cx) and the mouse tumors serially imaged by HFUS at the indicated times post-surgery. A. Three-dimensional prostate tumor reconstructions. HFUS images from the castrated mice (P364, P377, P380 and P386) were segmented in three dimensions with AMIRA software and the tumor pseudo-colored green. Tumors arise in both anterior lobes and HFUS images were captured separately for the right (R) and left (L) lobes. **B**. Kinetics of tumor volume following castration. Tumor volumes were determined from the 3D reconstructions, at the indicated times post-castration (n=4, blue) or post-sham operation (n=3, red) and plotted as percent pre-castration volume (mean [%] ± SEM).

**Figure 2.**
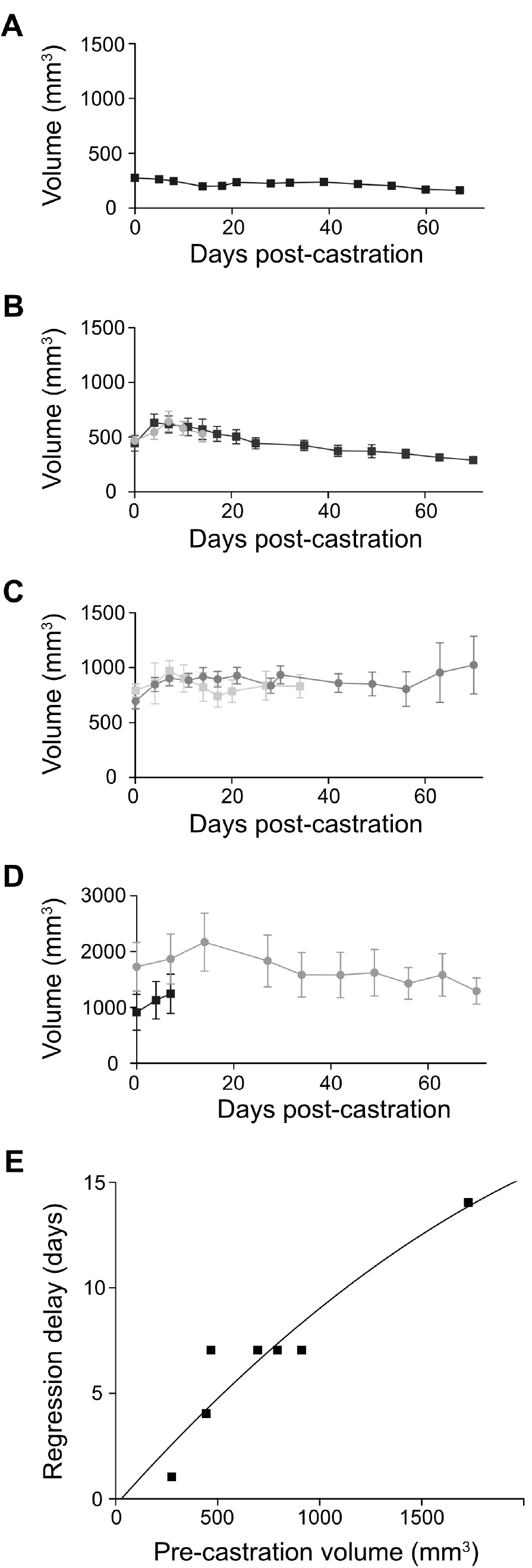
Regression delay is dependent on pre-castration tumor volume. Tumor-bearing PB-Cre4:Pten^*fl/fl*^ mice were castrated, tumors were serially imaged and volumes were determined using HFUS, followed by segmentation with AMIRA software, as described in Fig. 1. **A-D**. Kinetics of tumor volume following castration. Mean tumor volume for groups of mice (n = 4-6) corresponding to pre-castration tumor volumes of: ∼250 mm^3^ (**A**); Two groups ∼500 mm^3^ (**B**); Two groups ∼700 mm^3^ (**C**) and ∼1000 mm^3^ (black line) or ∼1700 mm^3^ (grey line) (**D**). For each panel. tumor volumes were determined from 3D reconstructions, at the indicated times post-castration and plotted as tumor volume (mean volume [mm^3^] ± SEM). **E**. Regression delay increases with pre-castration tumor volume. Line fit logarithmically, R^2^ = 0.878. Data derived from the groups shown in **A-D**.

To determine if the increase in tumor volume in castrated mice is due to an inflammatory infiltrate or accompanying edema, we castrated mice, sacrificed these animals at 6 and 14 days post-castration, fixed the tumors and examined H&E stained tissue sections. Sham operated animals provided controls. Pathological examination of the histological tissue sections revealed no significant inflammatory infiltration within the glands or within the surrounding stroma (Fig. 3A). Tissue sections from the same set of mice were immunohistochemically stained for Ki67 expression (Fig. 3B), to determine the rate of proliferation of the epithelial tumor cells within the prostate. When we ranked tumors by a Ki67 index (Fig. 3C), three (of the four) mice sacrificed at 6d post-castration were among the animals with the highest Ki67 indices observed in the sections from tumors of sham, 6d or 14d castrated mice. This suggests that there was a transient increase in the proliferation of the prostate tumor cells during the six days following castration which contributes to the volume increase that defines the delayed regression phenotype we have observed. Intra-tumoral androgen production could account for the transient proliferative response to castration that leads to the delay in regression. We therefore measured androgen levels in prostate tumor tissue. Following castration, testosterone and dihydrotestosterone levels decreased to undetectable levels and remained undetectable for at least two weeks (Fig. 3D-E). Other androgen precursors were not induced following castration (Supplementary Fig. S2B-D).

**Figure 3.**
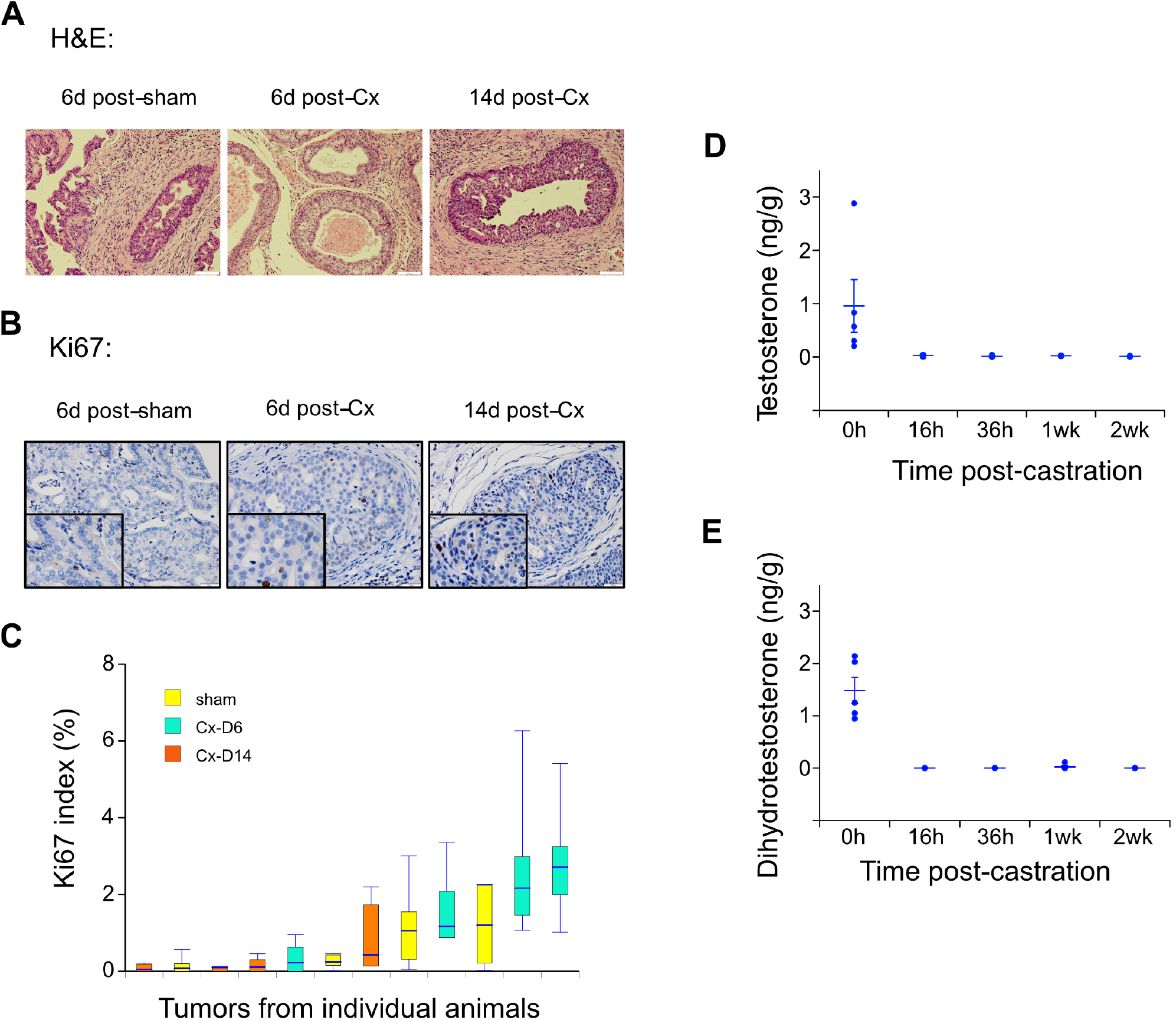
Castration reduces intratumoral androgens, and transiently increases tumor cell proliferation but does not induce an inflammatory cell infiltrate. Tumor-bearing PB-Cre4:Pten^*fl/fl*^ mice were sham-operated (sham, 0h) or castrated (Cx), sacrificed at the indicated times post-surgery. **A**. H&E staining of paraffin-embedded tumor tissue sections, 200x magnification. **B**. Immunohistochemical staining (brown) for Ki67, 400x magnification. **C**. Ranked-order box plot of Ki67 index for the indicated sets of tumors (n=4). The Ki67 index data were log transformed, plotted in rank-order and analyzed using a linear mixed model (F=20.05,2,70). **E-F**. Androgen levels were determined using LC/MS/MS from frozen tumor tissue (n=5 for 0h, 1wk, 2wk; n=4 for 18h, 36h). Levels were below the limits of detection (as described in **Methods)** after castration for 6/18 testosterone determinations and 17/18 DHT determinations. Tumor testosterone (**E**) and dihydrotestosterone (**F**) levels for individual mice, at the indicated times post-castration.

### Castration-induced regression delay is mediated by GR signaling

Since it appears that androgen-dependent signaling is not responsible for the regression delay we have observed, we hypothesized that glucocorticoid-GR signaling could partially compensate for the loss of AR signaling and mediate the regression delay. To test our hypothesis, we measured intra-tumoral endogenous ligands of GR. Two glucocorticoids that are active GR ligands (Brookes et al., 2012) – corticosterone and deoxycorticosterone – showed a modest increase post-castration (Fig. 4A). Interestingly, the levels of two precursors (5a-dihydroprogesterone and pregnenolone; Fig. 4B) were reduced, suggesting that sustained levels of corticosterone and deoxycorticosterone might be caused in large part by the generation of corticosterone from inactive 11-dehydrocorticosterone (Fig. 4B) by 11ß-HSD1, in the absence of 11ß-HSD2 (Li et al., 2017). ADT may also upregulate hexose-6-phosphate dehydrogenase, which generates NADPH required for 11ß-HSD1 activity (Li et al., 2021). Since the levels of glucocorticoid ligands were sustained, we sought to determine if GR signaling was necessary for the observed delay in regression, by inhibiting GR ligand binding using CORT125281. CORT125281 is one of a series of novel glucocorticoid receptor antagonists (Hunt et al., 2017; Morgan et al., 2002) that are more selective than older antagonists such as RU486 (Kach et al., 2017). Fig. 5B shows that treatment with CORT125281 (GRi) reduced the regression delay in PTEN-deficient tumors. Next, we examined these phenomena in a second murine model of primary prostate cancer (the Hi-MYC GEMM), driven by prostate epithelial cell-specific over-expression of c-MYC (Ellwood-Yen et al., 2003). We observed that castration of these mice was also followed by a delay in regression (Fig. 5A) with a remarkably similar kinetic profile to the PTEN-deficient model (compare Figs. 1B and 5A), although the delay appears to be closer to 3 days than 7 days. Importantly, administration of CORT125281 was also effective in reducing the delay in the onset of castration-induced regression in the Hi-MYC GEMM (Fig. 5B).

**Figure 4.**
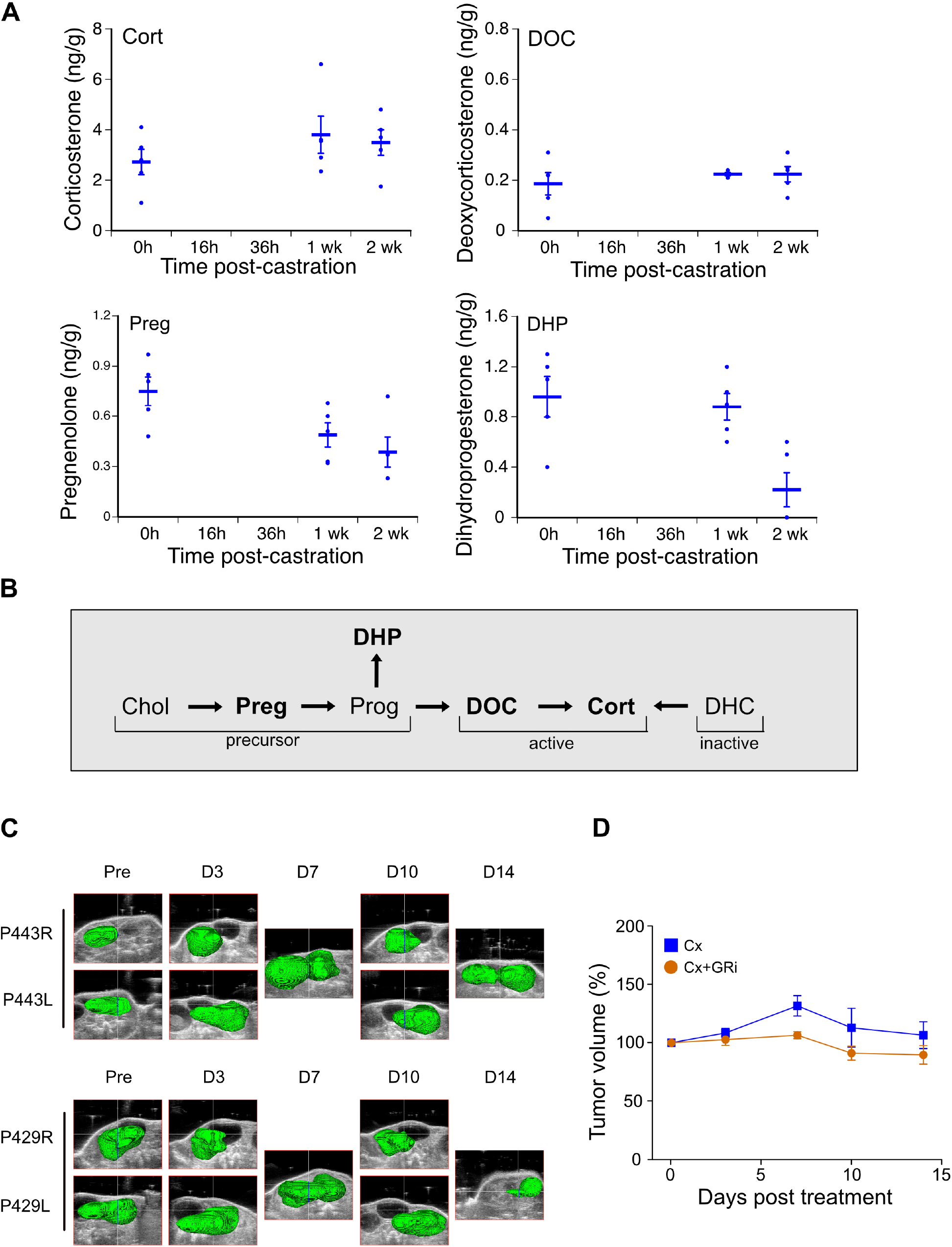
Inhibition of enhanced glucocorticoid signaling blocks castration-induced delayed regression in PTEN-deficient prostate cancers. **A**. Tumor glucocorticoid levels. Levels of the indicated glucocorticoids in three groups of tumor-bearing PB-Cre4:Pten^*fl/fl*^ mice from Fig. 4 were determined using LC/MS/MS. Levels of DHP were below the limit of detection for three 2wk samples. **B**. Scheme illustrating glucocorticoid steroid metabolism. Levels for steroids in bold type are shown in panel **A**. The arrows are not meant to reflect chemical equilibria but rather net synthesis. Steroid abbreviations (trivial name): Chol (cholesterol); Preg (pregnenolone); Prog (progesterone); DHP (5a-dihydroprogesterone); DOC (11-deoxycorti-costerone); Cort (corticosterone); DHC (11-dehydrocorticosterone). **C**. Tumor-bearing PB-Cre4:Pten^*fl/fl*^ mice were castrated only (Cx) or castrated and treated with CORT125281 (Cx+GRi). Tumors were serially imaged and volumes determined as in Fig. 1. Three-dimensional prostate tumor reconstructions from representative mice castrated and treated with CORT125281 (P443, P429) are shown. **D**. Kinetics of tumor volume following castration, in the presence of a GR inhibitor. Tumor volumes were determined from the 3D reconstructions, at the indicated times post-castration in mice receiving CORT125281 (n=5, orange) and plotted as percent pre-castration volume (mean [%] ± SEM). Tumor volumes from mice which were castrated and not treated are re-plotted from Fig. 1 (n=4, blue). Tumor volumes of each mouse are depicted in Supplementary Fig. S3.

**Figure 5.**
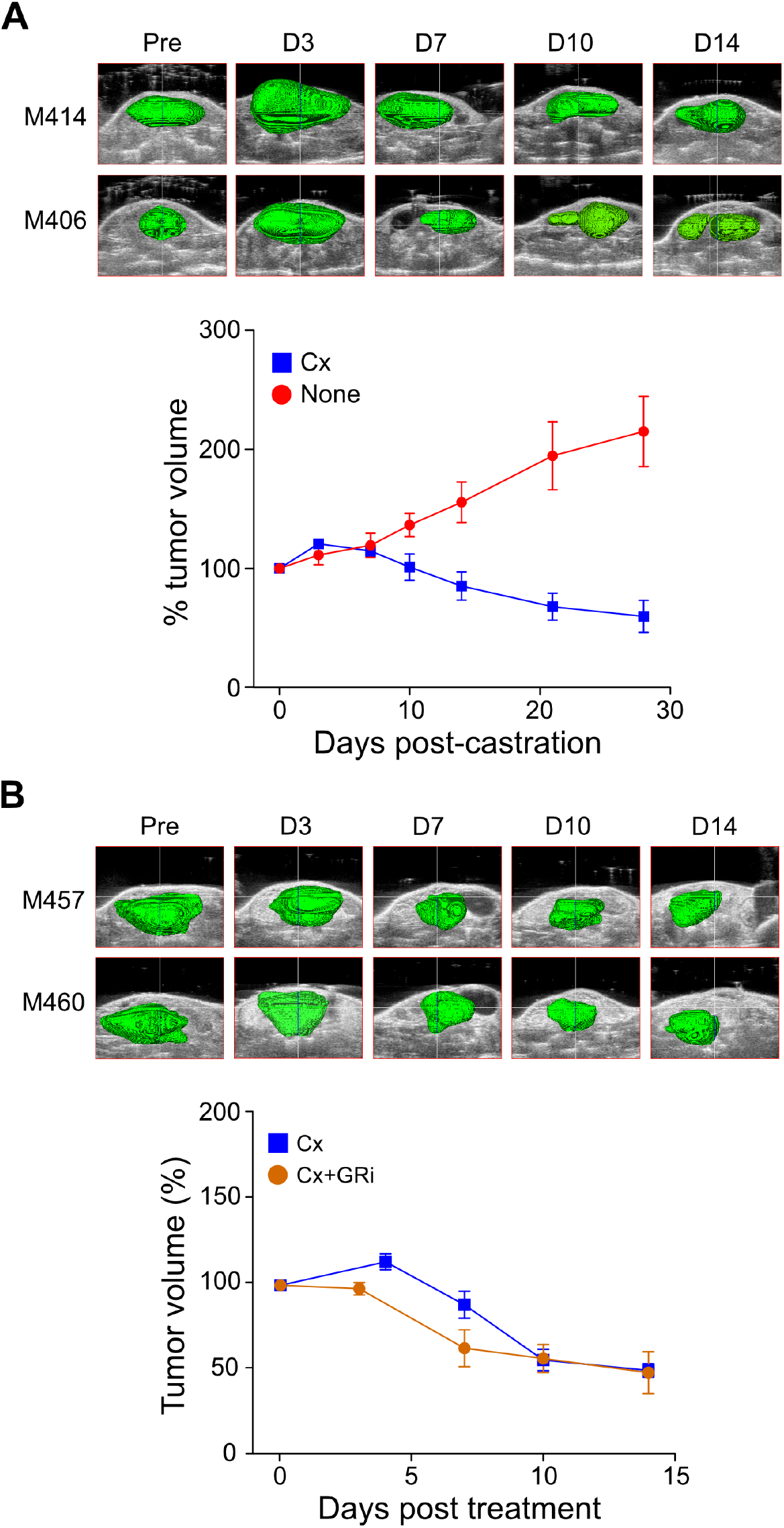
Castration-induced delayed regression of Hi-MYC prostate cancers is also dependent on glucocorticoid signaling. Tumor-bearing ARR_2_/PB-*Myc* mice were sham-operated (None), castrated (Cx) or castrated and treated with CORT125281 (Cx+GRi). Tumors were serially imaged and volumes determined as in Fig. 1. **A**. <u>Upper panel:</u> Three-dimensional prostate tumor reconstructions from representative castrated Hi-MYC mice (P414, P406). <u>Lower panel:</u> Kinetics of tumor volume following castration. Tumor volumes were determined from the 3D reconstructions, at the indicated times post-surgery in castrated (n=7, blue) or sham-operated (n=7, red) mice and plotted as percent pre-castration volume (mean [%] ± SEM). **B**. <u>Upper panel:</u> Three-dimensional prostate tumor reconstructions from representative Hi-MYC mice which were castrated and received CORT125281 (P457, P460). <u>Lower panel:</u> Kinetics of tumor volume following castration. Tumor volumes were determined from the 3D reconstructions, at the indicated times post-castration in mice receiving CORT125281 (n=5, orange) or mice which were only castrated and received no drug (n=6, blue). These values were plotted as percent pre-castration volume (mean [%] ± SEM). Tumor volumes of each mouse are depicted in Supplementary Fig. S3.

### GR expression and signaling is increased during the period of delayed regression

Since castration-induced enhanced GR signaling is required at least in part for the delayed regression phenotype (Fig. 4D, 5B), we measured changes in the level of GR protein and mRNA expression in tumor tissue, at the end of the delay period and compared those expression levels to the non-castrated tumor-bearing mice. Immunoblot analysis of prostate tumor lysates from both the PTEN-loss and MYC-gain GEMMs showed that near the end of the delay period (i.e., 3-4 days post-castration) the relative level of GR protein was elevated in both models (Fig. 6).

**Figure 6.**
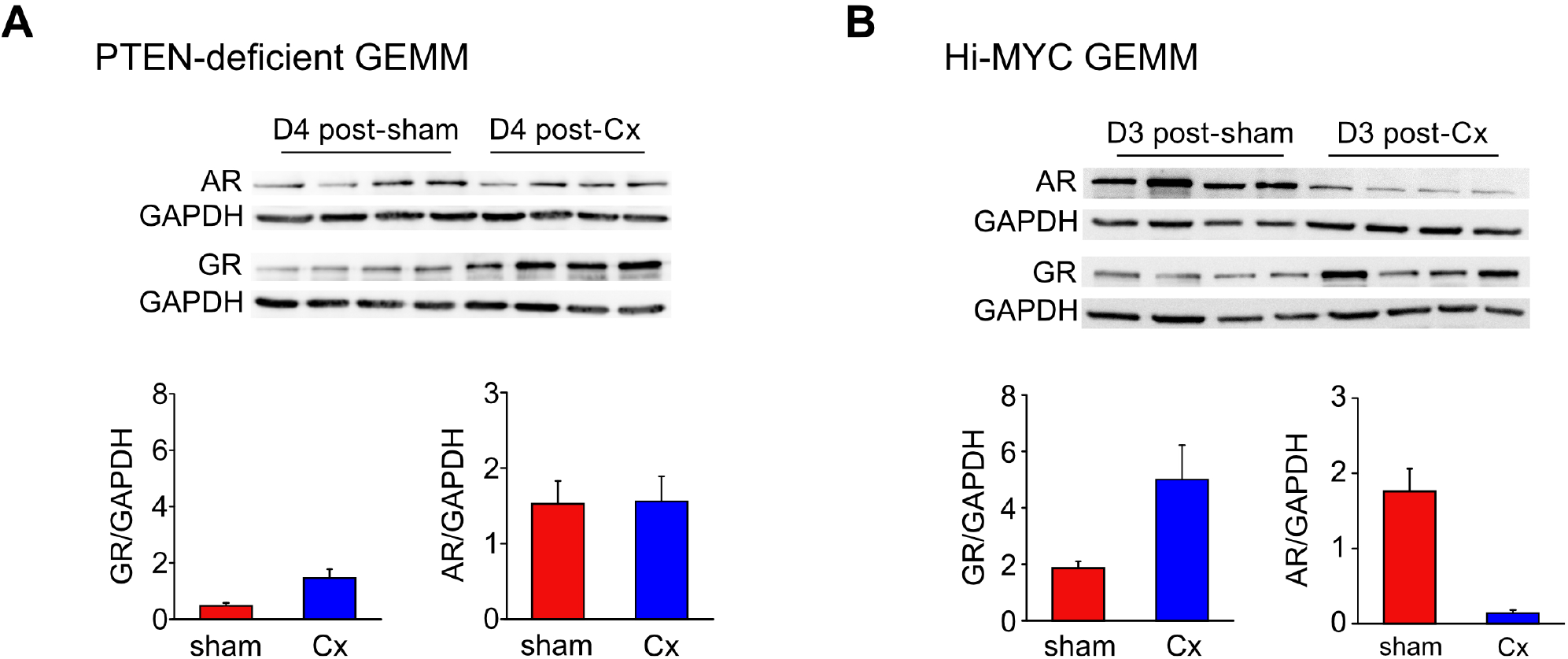
GR protein levels increase four days following castration. Groups (n=4) of tumor-bearing mice were sham-operated (SH) or castrated (Cx). Three or four days later (as indicated), mice were sacrificed, tumor tissue excised, protein lysates were prepared, immunoblot analyses of AR and GR proteins were performed in triplicate and protein levels were quantitated, as described in the **Materials and Methods. A**. PTEN-deficient GEMM. Representative immunoblots from tumors that developed in PB-Cre4 *P Pten*^*fl/fl*^ mice (upper panels) are shown. Relative levels of AR and GR protein, normalized to GAPDH, were derived from triplicate immunoblots (lower panels). **B**. Hi-MYC GEMM. Representative immunoblots from tumors that developed in ARR_2_/PB-*Myc* mice (upper panels) are shown. Relative levels of AR and GR protein, normalized to GAPDH, were derived from triplicate immunoblots (lower panels). Corresponding complete immunoblot images for both **A** and **B** are shown in Supplementary Fig. S4.

Next, we employed single cell RNA sequencing (scRNAseq) to assess changes in the GR mRNA, in the tumor cell population. Specifically, mice bearing PTEN-deficient tumors were either castrated (n=2) or sham operated (n=2), and sacrificed near the end of the delay period. Tumors were dissociated into single cells and these were subjected to RNA sequencing. Transcript counts from single cells of tumors from all four mice were pooled and the dimensionality of this transcriptomic data was reduced using the t-distributed stochastic neighbor embedding (t-SNE) algorithm. t-SNE visualization of the compiled transcript dataset, followed by hierarchical clustering and subjective cell annotation based on transcript function, allowed us to identify seven relevant cell types within the tumor population (Fig. 7A). In Fig. 7B, we projected CD45 (hematopoietic/immune cell lineage) and EPCAM (epithelial cell lineage) expression on to the t-SNE plot from Fig. 7A. This confirms the expected lineage of the putative cell population assignments. We then repeated the t-SNE analysis and hierarchical clustering for each of the four individual samples and determined the number of GR transcripts in both the entire GR transcript-positive cell population (all GR+ cells) and in the GR+ epithelial cell population, as shown in Supplementary Fig. S5 (upper panels). Pooling of like samples revealed that GR transcript levels are increased – four days post-castration, relative to sham-operated mice – in the epithelial cell population of the tumor (Fig. 7C), but not in the entire tumor cell population, which included stromal and immune cell populations, but not seminal vesicle (Supplementary Fig. S5, lower panels).

**Figure 7.**
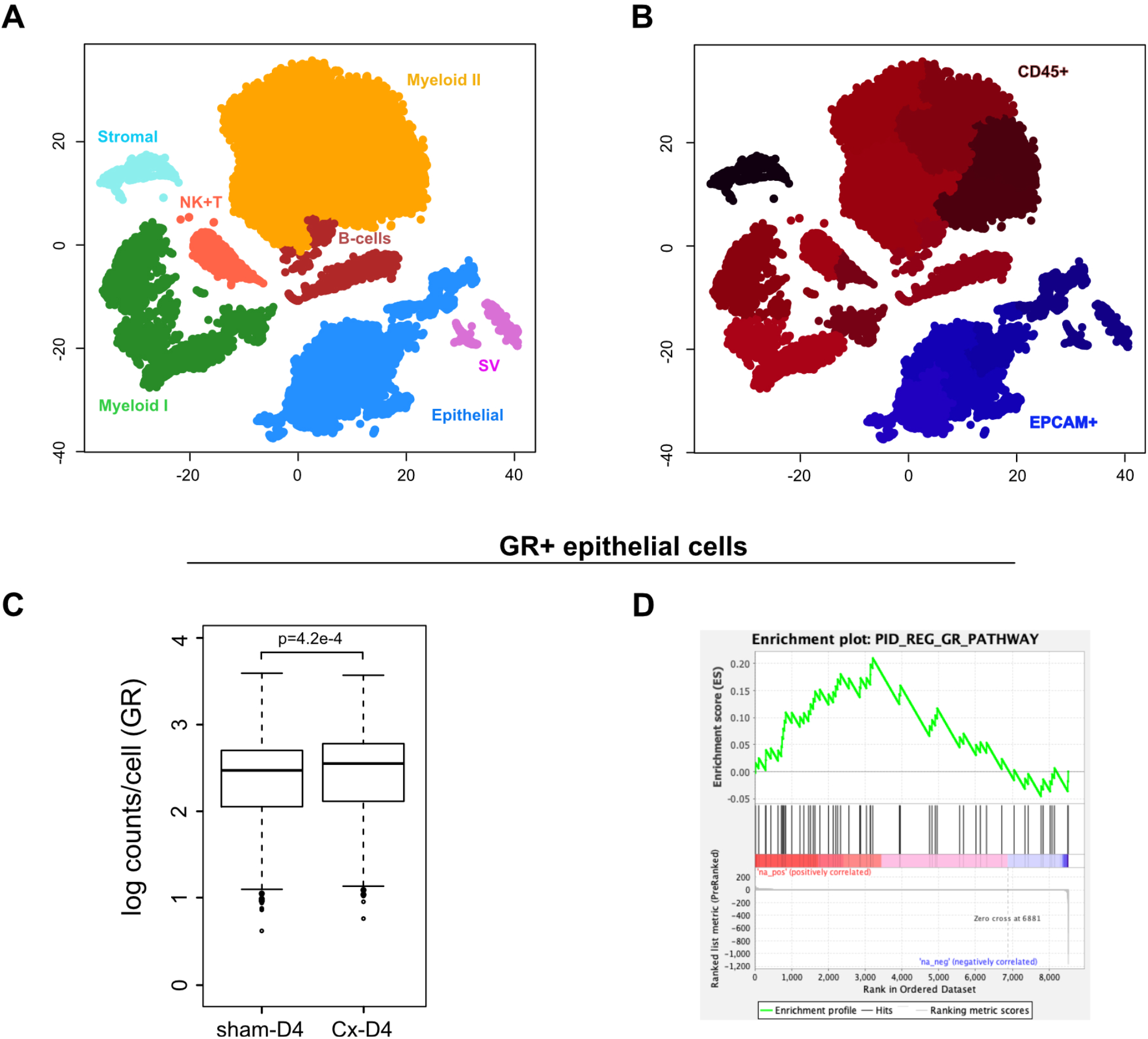
Single cell transcriptome analysis demonstrates that GR mRNA and GR-regulated genes are increased during the regression delay. Tumor-bearing PB-Cre4:Pten^*fl/fl*^ mice were castrated (n=2) or sham operated (n=2), sacrificed four days later and tumor tissue excised. The tumor was dissociated into single cells and for each sample RNA from about 5,000 cells was sequenced, as described in the **Materials and Methods. A**. t-Distributed stochastic neighborhood embedding (t-SNE) visualization of single cell mRNA data from all four mice, followed by hierarchical clustering and gene transcript annotation. Seven cell sub-populations were identified and color-coded as indicated by the labels in the panel: epithelial (tumor epithelial cells); NK+T (NK cells, T-cells); myeloid I and II (two distinct myeloid cell populations); stromal (fibroblast, myofibroblast and endothelial cells); B-cells; and SV (seminal vesicle). **B**. Relative expression of CD45 (red) and EPCAM (blue) mRNAs in the cell clusters identified in panel A. Darker color indicates higher level of mRNA expression (transcript counts). The stromal cluster is shaded black to indicate that it does not have significant expression of either mRNA. **C**. GR mRNA expression is increased four days post-castration (Cx-D4) relative to sham-operated mice (sham-D4). Box plot of transcript counts corresponding to GR-expressing epithelial cells identified in panel **A**. Data from individual mice is shown in Supplementary Fig. S5 (right side). **D**. GSEA demonstrates increased GR signaling during the regression delay. Differential gene expression of the pooled transcripts from the epithelial cell population followed by GSEA identified one glucocorticoid signaling gene set (PID_REG_GR_PATHWAY) that was significantly upregulated in the epithelial cell population.

Focusing on only those transcripts expressed by the epithelial cell cluster, we performed a differential gene expression analysis comparing the pooled transcript data from the prostate cancers of the sham-operated mice with those from the four-day post-castration mice. Gene-set enrichment analysis (GSEA) (Subramanian et al., 2005) of differentially expressed genes demonstrated that a glucocorticoid signaling gene set (PID_REG_GR_PATHWAY) was significantly upregulated in the epithelial cell population (Fig. 7D and Supplementary Fig. S6B). Importantly, in addition to the GR (N3RC1) itself, ∼50% (16/31) of the enriched genes are related to cell cycle, apoptosis, mitogenesis or tumorigenesis (Supplementary Table S1). In contrast, GSEA of the total tumor cell population – as in Fig. 7C – demonstrated that a separate glucocorticoid signaling pathway (WR_GLUCOCORTICOID_RECEPTOR_PATHWAY) was significantly upregulated (Supplementary Fig. S6A). Only ∼20% (3/14) of the enriched genes (plus GR/N3RC1) in this pathway are related to cell cycle, apoptosis, mitogenesis or tumorigenesis (Supplementary Table S2).

## DISCUSSION

We used HFUS imaging to determine the kinetics of tumor volume following castration in two murine models of prostate cancer driven by genetic alterations (PTEN loss, MYC over-expression) that are common in human prostate cancer. Surprisingly, we found that in the 4-7 day period immediately following castration-induced androgen deprivation, the prostate cancer tumor volumes in both models continued to increase, at a rate that was indistinguishable from tumors in untreated mice (Figs. 1 and 6). We monitored the kinetics of tumor volume following castration in mice grouped according to initial (pre-castration) tumor volume and found that the regression delay was proportional to the initial tumor volume (Fig. 2). During the regression delay there is an increase in tumor volume of 10-20%, yet there is no discernable inflammatory or immune cell infiltrate in H&E tissue sections taken from tumors harvested just prior to the beginning of tumor regression (Fig. 3A). Moreover, there is an increase in expression of the proliferative marker Ki67, detectable in the epithelial tumor cell population by the end of the delay period (Fig. 3B-C). Taken together, these data suggest that the unexpected increase in tumor volume following castration is due to continued proliferation of the epithelial tumor cell population, even as castration triggers the apoptotic process that ultimately leads to regression of the prostate cancer.

The studies of Huggins in the 1940’s first demonstrated that prostate cancer growth is dependent on testicular androgens (Huggins and Hodges, 1941). In the 80 years since, it has become clear that proliferation is dependent on signaling through the androgen receptor. In prostate cancer, a variety of initiating mutations in FoxA1, HoxB13, SPOP and other genes converge on a common mechanism – reprogramming of the AR cistrome – inducing the transcription of a set of genes that drive tumor cell proliferation in response to androgens (Copeland et al., 2019; Grbesa et al., 2021; Pomerantz et al., 2015). Since we did not detect any residual androgens in tumor extracts from the PTEN-deficient mice (Fig. 3), we surmised that conventional signaling through the AR has been turned off. Therefore, we sought to determine whether signaling through the GR, a member of the nuclear receptor super-family that is closely related to the AR, substitutes for AR signaling following castration and during the subsequent delayed regression period. Indeed, our key finding is that CORT125281, a highly selective glucocorticoid receptor antagonist (Hunt et al., 2017; Morgan et al., 2002), reduced the regression delay in both the PTEN-deficient (Fig. 4D) and the Hi-MYC (Fig. 5B) prostate cancer models. This strongly supports a role for GR-signaling in the delay period that follows castration.

We examined the PTEN-deficient model in detail at the molecular level and, consistent with the requirement for GR signaling, we observed an increase in key components of the signaling pathway: glucocorticoid ligands (Fig. 4A) as well as the glucocorticoid receptor (both protein and mRNA; Figs. 6A and 7C, respectively). We used single cell transcriptomic analysis to determine the changes in the level of GR mRNA in the epithelial tumor cell population only, since GR is widely expressed and serves different functions in different cell populations within the tumor and its associated microenvironment. Indeed, when we combined the scRNAseq data from all of the cell populations we identified in Fig. 7A, we did not observe a difference in the number of GR transcripts per cell (Supplementary Fig. S5). When we performed bulk RNAseq analysis on similar tumors, we also saw no difference in GR transcripts/cell at the same time points post-castration (data not shown) further confirming the utility and resolving power of the single cell methodology. Another important observation from our transcriptomic analysis concerns the GR-regulated gene sets identified by GSEA. When we analyzed the epithelial cell population only, we found a gene set that had a larger fraction of genes which we subjectively identified as having a tumorigenic or proliferative function, relative to the GR gene set identified by GSEA of all of the cell populations depicted in the t-SNE plot (Supplementary Tables S1-S2). Finally, although we did not perform a complete set of molecular analyses on the Hi-MYC model, we observed that GR protein was also elevated in tumors post-castration (Fig. 6B), similar to our observations in the PTEN-deficient GEMM (Fig. 6A). This suggests a similar mechanism is operative in two genetically distinct models.

Similar studies from Xie *et al*. (Xie et al., 2015) strongly support our observations. First, they describe a transcriptional model where AR signaling negatively regulates GR/N3RC1 transcription. This mechanism may well be operative in our GEMMs, although it does not account for the transient nature of the regression delay that we have documented. Second, in a model of ADT that employs AR gene silencing, there is a clear pattern of delayed regression that is very similar to the pattern we observe in Figs. 1, 2 and 6A, although these authors do not comment on this aspect of their data (Xie et al., 2015). Finally, they demonstrate that GR expression is induced in samples of human prostate cancer, when patients are given neo-adjuvant hormonal therapy (Xie et al., 2015).

The immediate, transient response to androgen that we have documented – an increase in GR signaling that is a consequence of both increased levels of active glucocorticoids and increased expression of the GR – is very similar to the response to AR signaling blockade that has been observed in a subset of castration resistant prostate cancers. Specifically, Arora *et al*. (Arora et al., 2013) and others (Isikbay et al., 2014; Puhr et al., 2018) have shown that GR substitutes for the AR to overcome enzalutamide resistance in murine models of prostate cancer and that the mechanism involves increased GR expression. Indeed, CORT125281, renamed relacorilant, is currently in early phase trials in combination with enzalutamide for patients suffering metastatic CRPC (NCT03674814, NCT03437941).

Sharifi and colleagues (Li et al., 2017; Li et al., 2021) have documented that sustained levels of glucocorticoids is another mechanism that can contribute to GR ‘takeover’ in enzalutamide resistant prostate cancers. In human cells, they have identified mechanisms that are driven by androgen suppression and lead to the production of cortisol from the inactive metabolite cortisone, by degrading 11ß-HSD2 (Li et al., 2017) or enhancing the reductive activity of 11ß-HSD1 via increased H6PD expression (Li et al., 2021). An analogous set of mechanisms could generate active corticosterone from inactive 11-dehydrocorticosterone (Fig. 4B) in our models, following castration.

Finally, our observation of sustained proliferation in the context of androgen deprivation, driven by GR signaling has potential implications for prostate cancer therapy, as others have noted (Arora et al., 2013; Isikbay et al., 2014; Valle and Sharifi, 2021). Residual GR signaling dependent lineages may support genetic instability leading to cells expressing AR variants or harboring AR amplification, or emergence of neuroendocrine/stem cell-like AR signaling independent lineages. Specifically, our data support the idea of combining AR and GR inhibition and also reducing the use of glucocorticoids in combination with other therapies for castration resistant prostate cancers.

## ACKNOWLEDGEMENTS

We thank Corcept Therapeutics for providing CORT125281. This work was supported by the Department of Defense Prostate Cancer Research Program No. W81XWH-19-1-0378 (to J.J. K), the S.A.S. Foundation and the ACS (to K.L.N), and by institutional funds from the Roswell Park Alliance Foundation and the National Cancer Institute Cancer Center Support Grant P30-CA016056 to Roswell Park Comprehensive Cancer Center.

## AUTHOR CONTRIBUTIONS

Conceptualization, K.L.N. and J.J.K.; Methodology, R.Z., K.H.E., K.L.N., and J.J.K.; Investigation, A.M., R.Z., K.S., S.S., and C.P.; Formal Analysis, B.X., M.M. and K.H.E. Writing – Original Draft, A.M., K.L.N., and J.J.K.; Writing – Review & Editing, B.X., G.C., K.H.E., K.L.N. and J.J.K.; Funding Acquisition, K.L.N. and J.J.K.; Supervision, K.L.N. and J.J.K.

## METHODS

### Key Resource Table

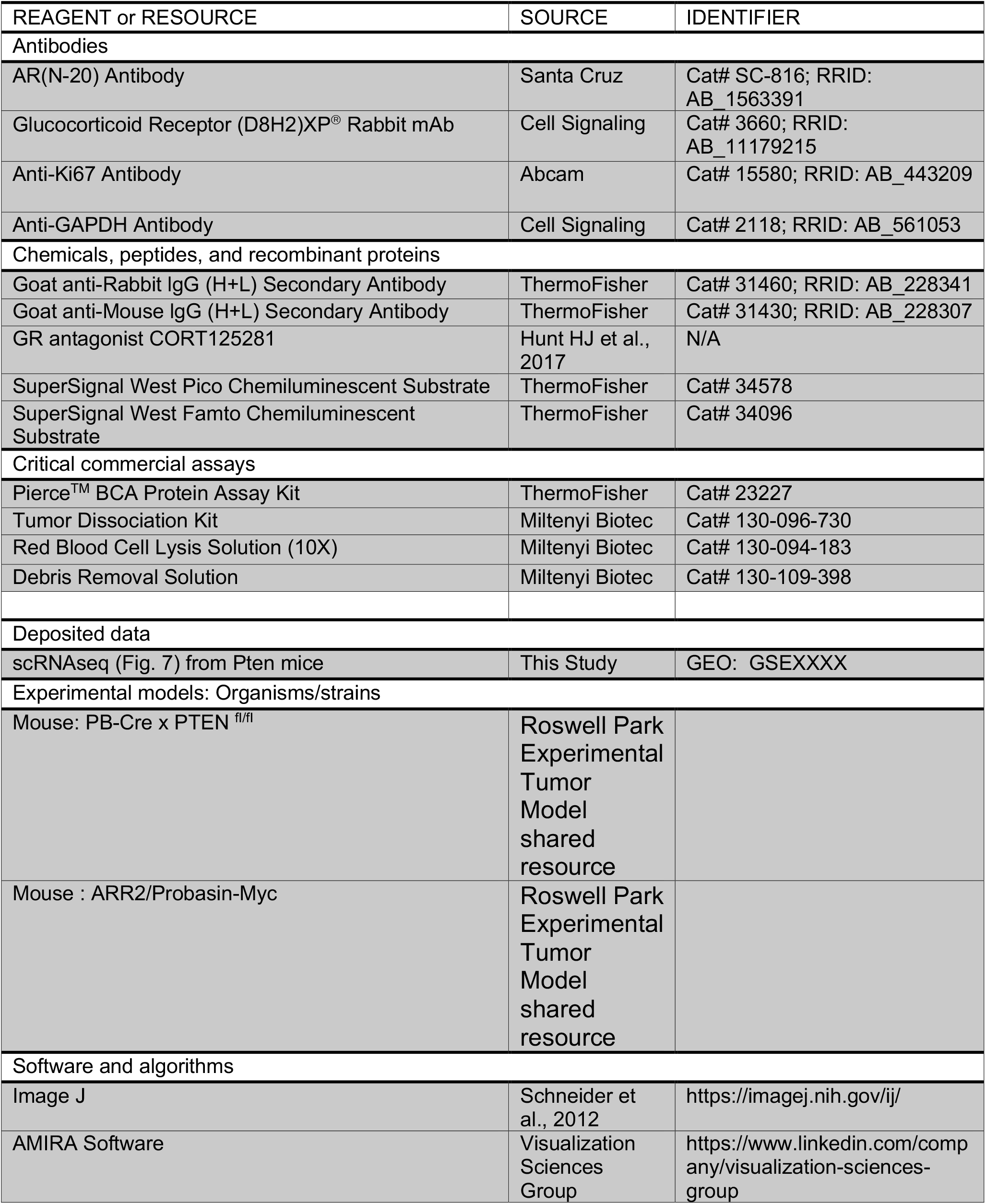
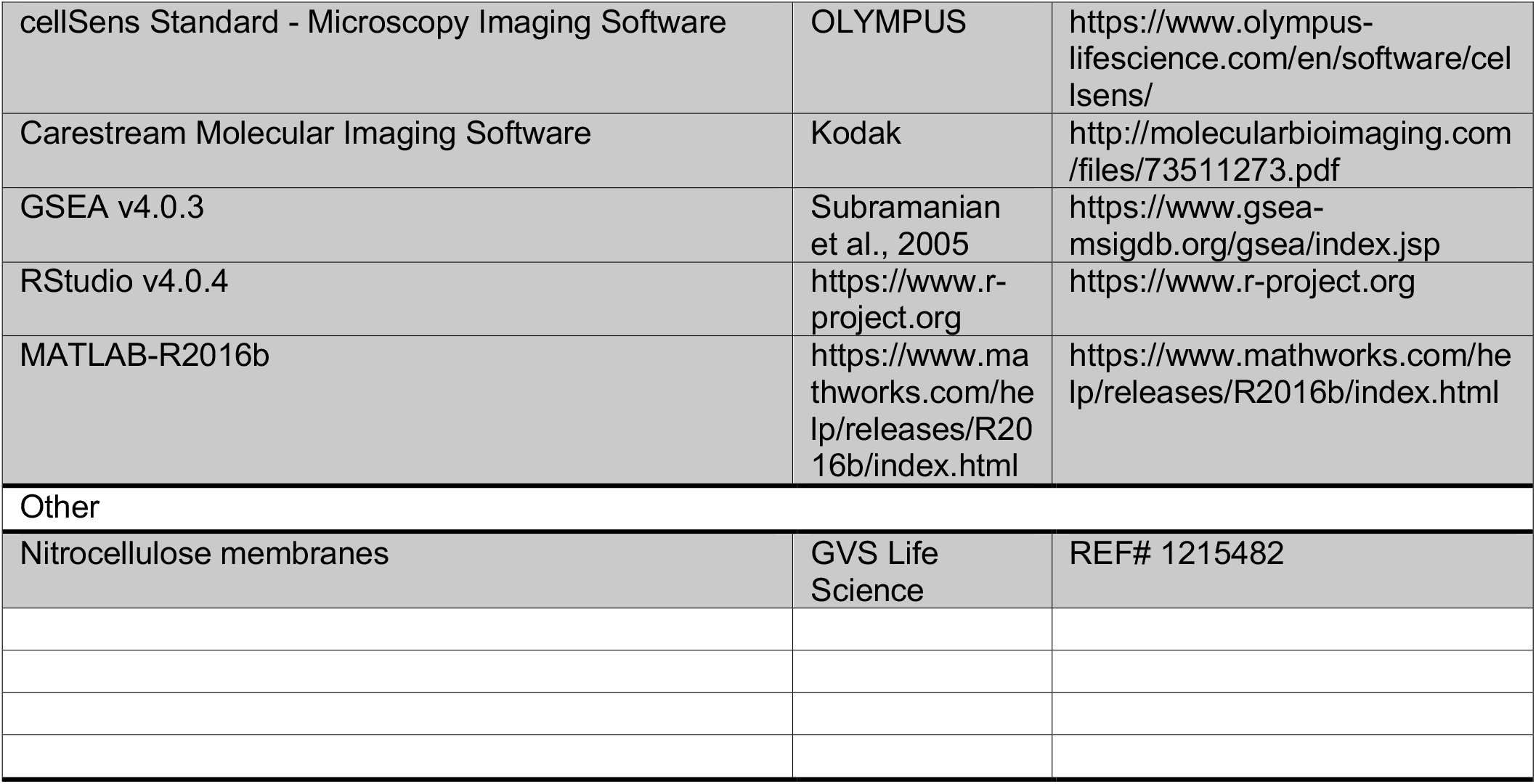

## RESOURCE AVAILABILITY

### Lead contact

Further information and requests for resources and reagents should be directed to and will be fulfilled by the Lead Contact, Kent L. Nastiuk

### Materials availability

This study did not generate new unique reagents. We received the compound CORT125134 through an MTA with Corcept Therapeutics.

### Data availability

The gene expression data are publicly available from the National Center for Biotechnology Information GEO (GSE). The remainder of the data generated in this study are available within the article and its supplementary data files.

## EXPERIMENTAL MODEL AND SUBJECT DETAILS

### Genetically engineered mouse prostate cancer models

Two genetically engineered mouse models of prostate cancer were bred in the Roswell Park Experimental Tumor Model shared resource, maintained on a 12-hour light, 12-hour dark cycle in a temperature and humidity controlled room with free access to food and water. PB-Cre4 x PTEN^fl/fl^ mice (Wang et al., 2003) were generated by crossing male PB-Cre4 (Frederick Lab, NCI) and female PTEN^fl/fl^ (Jackson Labs) mice. Transgenic ARR_2_/probasin-*Myc* (Hi-MYC) mice (Ellwood-Yen et al., 2003) were from Frederick Lab, NCI. For both GEMMs, male pups were genotyped and monitored for tumor development by imaging (see below) beginning at 12 weeks. Mice were enrolled in experiments when tumors were >300 mm^3^ and animals were older than 6 months age. Castration and imaging were performed using isoflurane anesthetic, as described previously (Singh et al., 2015). In some cases, mice received 20mg/kg/day intraperitoneal injections of CORT125281 (Koorneef et al., 2020), which was reconstituted at 40mg/ml in DMSO and further diluted with sesame oil. At sacrifice, mouse prostate tumors were identified grossly, excised and bisected; half frozen in liquid nitrogen and half rinsed and fixed in formaldehyde. All animal studies were performed in accordance with the National Institute of Health Guidelines for the Care and Use of Laboratory Animals and Use of Laboratory Animals and approved by Roswell Park Institutional Animal Care and Use Committee (1308M).

## METHOD DETAILS

### Three-dimensional ultrasound imaging

Ultrasound imaging of prostate tumors was performed using a Vevo 2100 micro-ultrasound imaging system (Vevo LAZR; VisualSonics Inc., Toronto) with a 21-MHz linear-array transducer system (LZ250; 13-24 MHz maximal broadband frequency; 21 MHz center frequency, 75 μm axial resolution, 80 μm lateral resolution, 23 mm maximal lateral field of view) in the Roswell Park Translational Imaging shared resource. Briefly, mice were anesthetized with isoflurane, the abdomen was depilated and ultrasound transmission gel was applied. Three-dimensional scans of ultrasound B-mode image being recorded digitally and 3D stacks of these B-mode images were imported into AMIRA software (Visualization Sciences Group, Burlington MA) for tumor volume reconstruction.

### Histology and immunohistochemistry

Mouse prostate tumor halves intended for histological analysis were fixed in 10% formaldehyde for two days, embedded in paraffin, and cut into 5 um sections onto glass slides. Individual sections were stained with hematoxylin and eosin (H&E) or immunohistochemically stained using an antibody to Ki67 (Abcam, cat. no. 15580; 1:1000), on an automated staining platform in the Roswell Park Experimental Tumor Model shared resource. Stained sections were viewed and photographed on an OLYMPUS BX45 microscope. In the case of Ki67 IHC, representative ROIs were identified and positive stained tumor cell nuclear were counted, and divided by the number of total tumor cell nuclei to generate a Ki67 index (% positive staining tumor nuclei) from 6 fields for each tumor.

### Immunoblot analysis

Protein extracts were prepared by grinding frozen tissue with a pestle in a mortar containing liquid nitrogen. The resulting frozen powder was dissolved in lysis buffer, sonicated for 8 seconds at 10 watts and insoluble protein and debris were pelleted by high-speed centrifugation. Extracts from PTEN-deficient mouse tumors were prepared in a 1% SDS lysis buffer (1% SDS, 20mM Tris-HCl pH7.5, 150mM NaCl, 1% NP-40, 1mM EDTA, 1mM PMSF, 1x Proteinase inhibitor cocktail (Millipore Sigma)). Extracts from Hi-MYC mouse tumors were prepared in RIPA lysis buffer (0.5% sodium deoxycholate, 0.1% SDS, 20mM Tris-HCl pH7.5, 150mM NaCl, 1% NP-40, 1mM EDTA, 1mM PMSF, 1x Protease inhibitor cocktail (Sigma). Protein concentration was determined using bicinchoninic acid reagent (Pierce, ThermoFisher Scientific). Immunoblotting was performed essentially as described previously (Pan et al., 2020), with the following modifications. Twenty-five to 40 µg of protein extract were separated on 7.5% SDS-PAGE. Primary antibodies and working dilutions were: anti-AR (Santa Cruz, cat. no. SC-816, 1:500); anti-GR (Cell Signaling, clone D8H2, 1:1000) and anti-GAPDH (Cell Signaling, cat. No. 2118, 1:2000). Specific protein signal was captured using a cooled charge-coupled digital camera system (Kodak 4000R), quantitated using Kodak Molecular Imaging software and normalized using GAPDH levels, as indicated.

### Steroid metabolite determinations

Steroid levels were determined in samples of powdered frozen tumor tissue by the Roswell Park Bioanalytics, Metabolomics and Pharmacokinetics shared resource, using LC/MS/MS methodology, as previously described (Wilton et al., 2014). Limits of detection are: ASD and T = 31.25 pg/g; DHT = 62.5 pg/g; DHEA, 5a-dione and AND = 1000 pg/g, Cort, Preg, DOC, DHP =∼125 pg/g.

### Single cell RNA-sequencing and data analysis

Mouse prostate tumor tissue was dissociated for single cell RNA sequencing using the Tumor Dissociation kit from Miltenyi Biotec (cat. no. 130-096-730). Briefly, tissue was excised, washed with chilled 1x PBS, placed in a petri dish on ice, cut into 2-4 mm^3^ pieces, transferred to a gentleMACS C tube containing DMEM/enzyme mix and then attached to gentleMACS Octo Dissociator with heating running the ‘37C_m_TDK_1’ program. The suspension was filtered through a 40 µm cell strainer (Falcon). The cells were harvested by centrifugation at 400g for 10 min and resuspended in chilled 1X Red Blood Cell Removal Solution (Miltenyi Biotec, cat. no.130-094-183). Debris was then removed using the Debris Removal Solution (Miltenyi Biotec, cat. no. 130-109-398) following the manufacturer’s instructions.

Single cell gene expression libraries were generated using the 10X Genomics Chromium platform and sequenced on a NovaSeq 6000 (Illumina). Count data from all cells (n = 26,011) were included in a ‘SingleCellExperiment’ (version 1.8.0) object (Amezquita et al., 2020). Quality control, analysis and visualization of scRNAseq data were performed in R (version 3.6.0). Transcripts from a given cell were removed from the analysis if the corresponding transcriptome met any of the following criteria: 20% or more reads aligned to mitochondrial genes, 1500 total counts per cell or less, or 600 genes per cell or less. This yielded a total of 18,166 cells for analysis. The ‘scater’ package (version 1.14.5) was used to normalize counts per million (CPM) (McCarthy et al., 2017). The dimensionality of this transcriptomic data was reduced using the t-distributed stochastic neighbor embedding (t-SNE) algorithm (van der Maaten and Hinton, 2008). The ‘limma’ package (Ritchie et al., 2015) was used to identify differentially expressed genes to refine the assignment of clusters generated by the t-SNE algorithm. Briefly, clusters with high-level expression of EPCAM, KRT8, KRT18 and KRT19 were identified as epithelial cells; B cell populations were determined by CD79A, CD79B, CD19; NK and T cells by NKG7, KLRD1, CD3D, CD3E and CD3G; myeloid I by H2 genes and Cd74; myeloid II by S100A8/A9 and cytokines including CCL3, CCL4; non-immune stromal cells by COL3A1, COL1A2, and CD34; and seminal vesicle epithelial (SV) cells by SVS2, SVS5 and KRT18. Only cells with non-zero counts of NR3C1 (GR) were used to compare GR mRNA expression. To determine if glucocorticoid-related gene transcription was induced post-castration, we conducted gene set enrichment analysis (GSEA) (Subramanian et al., 2005) for ‘all cells’ (seminal vesicle excluded) and ‘epithelial (tumor) cells’, respectively. Low variance, low expression genes of all cells and epithelial cells were removed before performing GSEA. GSEA software (version 4.1.0) was used with classic enrichment, inclusion gene set size between 15 and 500, and the number of permutations set to 1000. A P-value cutoff of 0.05 with a false discovery rate (FDR q-val) cutoff of 0.25 were considered statistically significant.

## SUPPLEMENTARY TABLES

**Supplementary Table 1:** Genes from PID_REG_GR_PATHWAY enriched in tumor epithelial cells.

| SYMBOL | RANK IN<br>GENE LIST | RANK METRIC<br>SCORE | RUNNING ES | CORE<br>ENRICHMENT |
| --- | --- | --- | --- | --- |
| STAT1 | 20 | 46.70607376 | 0.016506929 | Yes |
| CDKN1A | 108 | 19.34492683 | 0.025104519 | Yes |
| EGR1 | 297 | 10.04787254 | 0.021779079 | Yes |
| IRF1 | 303 | 9.985536575 | 0.040056754 | Yes |
| FOS | 438 | 8.094637871 | 0.043106005 | Yes |
| NR4A1 | 633 | 6.239775658 | 0.039072264 | Yes |
| JUN | 724 | 5.765926361 | 0.047315706 | Yes |
| MAPK14 | 743 | 5.653651714 | 0.064058736 | Yes |
| GATA3 | 773 | 5.489055157 | 0.079503216 | Yes |
| NR3C1 | 807 | 5.352002144 | 0.09447549 | Yes |
| ICAM1 | 842 | 5.218481064 | 0.10932973 | Yes |
| SFN | 1006 | 4.534712791 | 0.10895553 | Yes |
| POU2F1 | 1229 | 3.750123978 | 0.1016164 | Yes |
| STAT5B | 1334 | 3.453756094 | 0.108207144 | Yes |
| RELA | 1463 | 3.156321526 | 0.11196469 | Yes |
| CREBBP | 1514 | 3.016171217 | 0.12493012 | Yes |
| KMT5B | 1591 | 2.888596058 | 0.13482626 | Yes |
| SMARCC1 | 1637 | 2.803568363 | 0.14838195 | Yes |
| VIPR1 | 1770 | 2.590506792 | 0.1516673 | Yes |
| MAPK8 | 2006 | 2.241907597 | 0.14279352 | Yes |
| AKT1 | 2131 | 2.077228785 | 0.14702326 | Yes |
| SMARCC2 | 2200 | 1.983715415 | 0.1578638 | Yes |
| PBX1 | 2251 | 1.92462647 | 0.17082924 | Yes |
| NCOA1 | 2331 | 1.827400446 | 0.18037122 | Yes |
| PRKACA | 2559 | 1.603734374 | 0.17244184 | Yes |
| EP300 | 2856 | 1.305983067 | 0.15636702 | Yes |
| CREB1 | 2877 | 1.294376493 | 0.17287394 | Yes |
| STAT5A | 3036 | 1.15143168 | 0.17309 | Yes |
| HSP90AA1 | 3142 | 1.076227903 | 0.1795627 | Yes |
| FKBP4 | 3145 | 1.075295925 | 0.19819452 | Yes |
| NCOA2 | 3206 | 1.041746736 | 0.20997946 | Yes |

Orange shading identifies genes associated with cell cycle, mitogenesis, apoptosis or tumor growth

**Supplementary Table 2:**
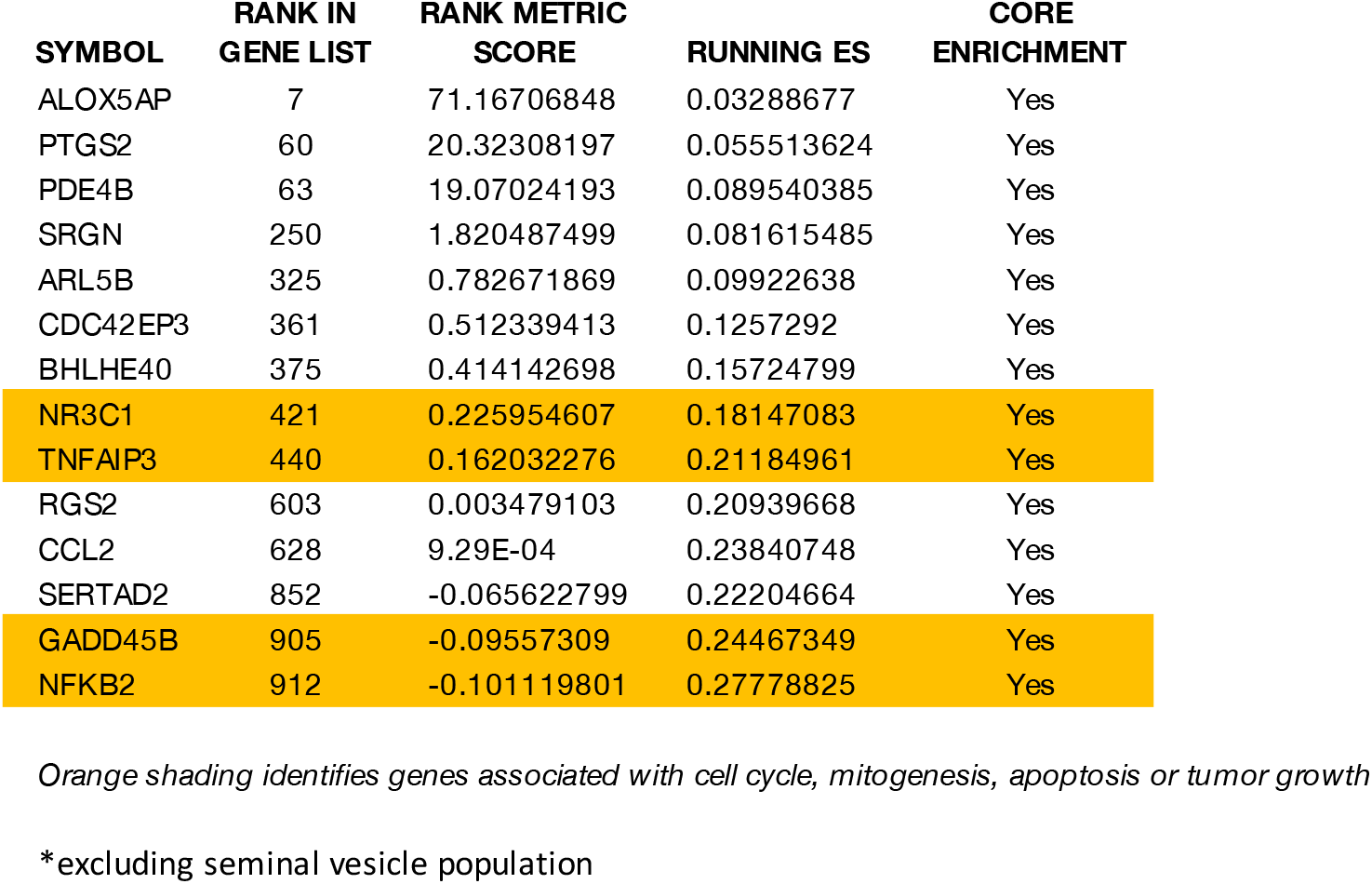
Genes from WP_GLUCOCORTICOID_RECEPTOR_PATHWAY enriched in all cells*.

| SYMBOL | RANK IN<br>GENE LIST | RANK METRIC<br>SCORE | RUNNING ES | CORE<br>ENRICHMENT |
| --- | --- | --- | --- | --- |
| ALOX5AP | 7 | 71.16706848 | 0.03288677 | Yes |
| PTGS2 | 60 | 20.32308197 | 0.055513624 | Yes |
| PDE4B | 63 | 19.07024193 | 0.089540385 | Yes |
| SRGN | 250 | 1.820487499 | 0.081615485 | Yes |
| ARL5B | 325 | 0.782671869 | 0.09922638 | Yes |
| CDC42EP3 | 361 | 0.512339413 | 0.1257292 | Yes |
| BHLHE40 | 375 | 0.414142698 | 0.15724799 | Yes |
| NR3C1 | 421 | 0.225954607 | 0.18147083 | Yes |
| TNFAIP3 | 440 | 0.162032276 | 0.21184961 | Yes |
| RGS2 | 603 | 0.003479103 | 0.20939668 | Yes |
| CCL2 | 628 | 9.29E-04 | 0.23840748 | Yes |
| SERTAD2 | 852 | -0.065622799 | 0.22204664 | Yes |
| GADD45B | 905 | -0.09557309 | 0.24467349 | Yes |
| NFKB2 | 912 | -0.101119801 | 0.27778825 | Yes |

*Orange shading identifies genes associated with cell cycle, mitogenesis, apoptosis or tumor growth*
\*excluding seminal vesicle population

## SUPPLEMENTARY FIGURES

**Figure S1.**
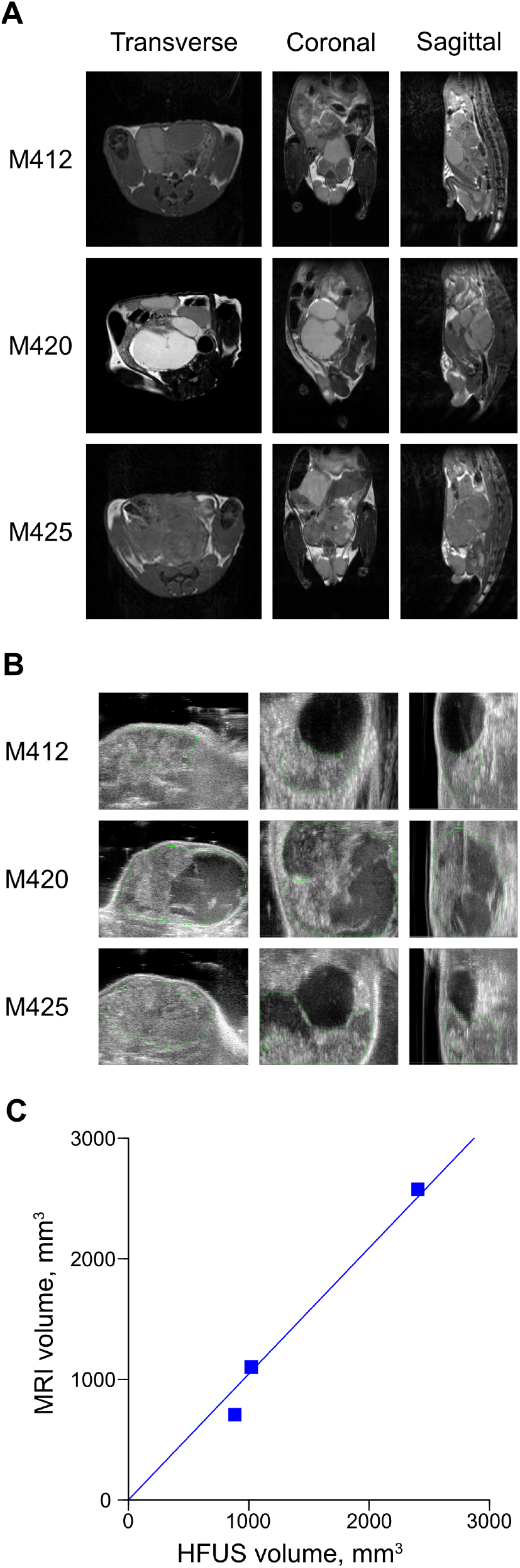
Comparison of MR and HFUS imaging protocols in Hi-MYC GEMM. Hi-MYC mice (n=3) harboring tumors of varying size were imaged using both MR and HFUS imaging protocols, as previously described (Nastiuk et al., 2007; Singh et al., 2015), to assess tumor volume. Representative images from MR (**A**) and HFUS (**B**) imaging protocols. **C**. Plot of tumor volumes determined by MR and HFUS imaging, demonstrating a high degree of correlation (r^2^ = 0.99).

**Figure S2.**
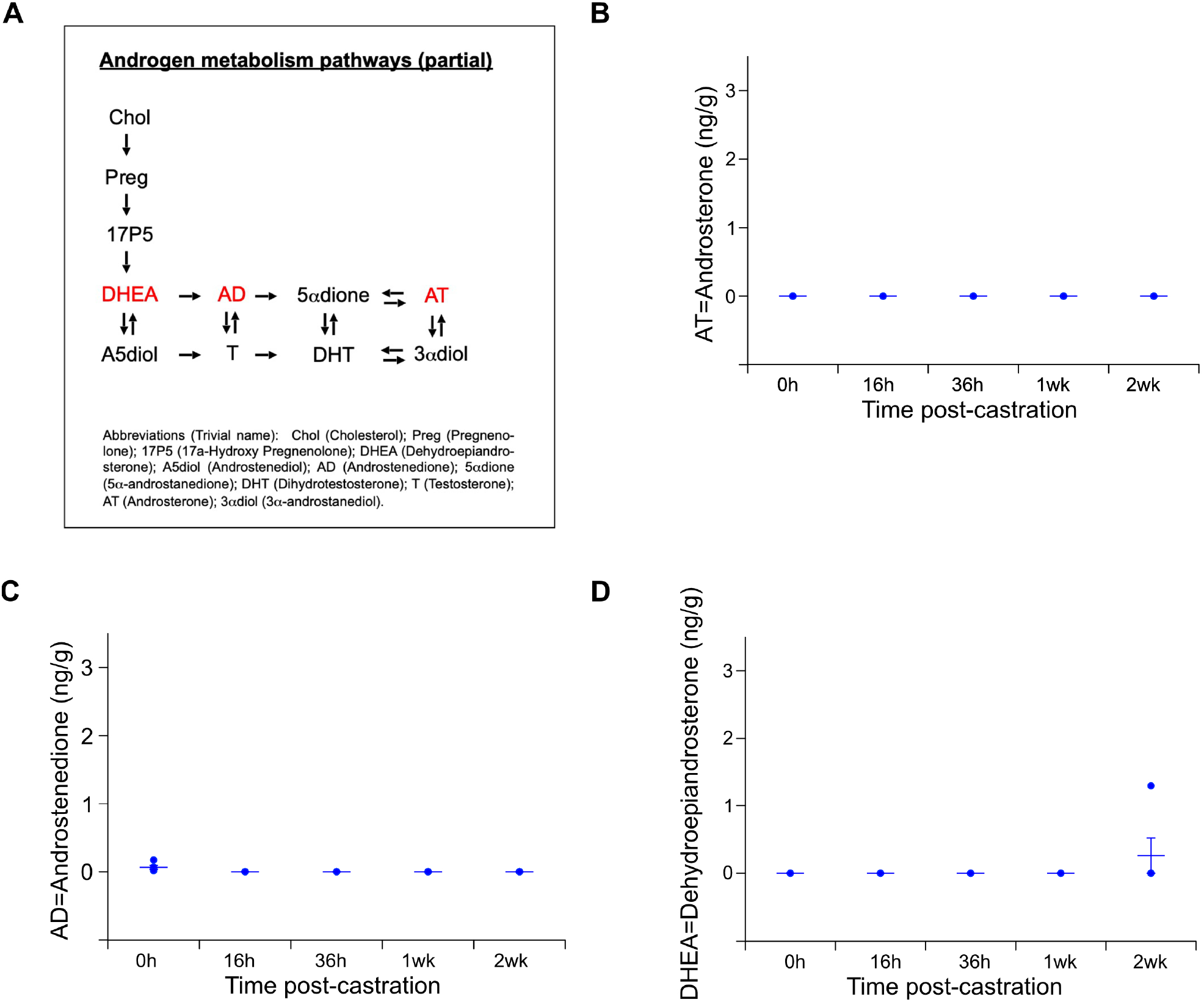
Intra-prostatic androstenedione (AD), dehydroepiandrosterone (DHEA) and androsterone (5α-dione) were undetectable in PTEN-deficient mice. Tissue samples from the mice employed in Fig. 4 were further analyzed for testosterone and dihydrotestosterone precursors (shown in red, in **A**), as described in the **Materials and Methods. A**. Partial scheme of androgen biosynthesis adapted from Sharifi (Sharifi, 2013). Abbreviations and the corresponding trivial names are shown in the inset. **B-D**, Levels of the androgen precursors androstenedione, androsterone and dehydroepiandrosterone were below the limits of detection in pre- and post-castration prostate cancer tissue samples, with the exception of 3 pre samples for androstenedione, and one 14d post-castration sample for DHEA.

**Figure S3.**
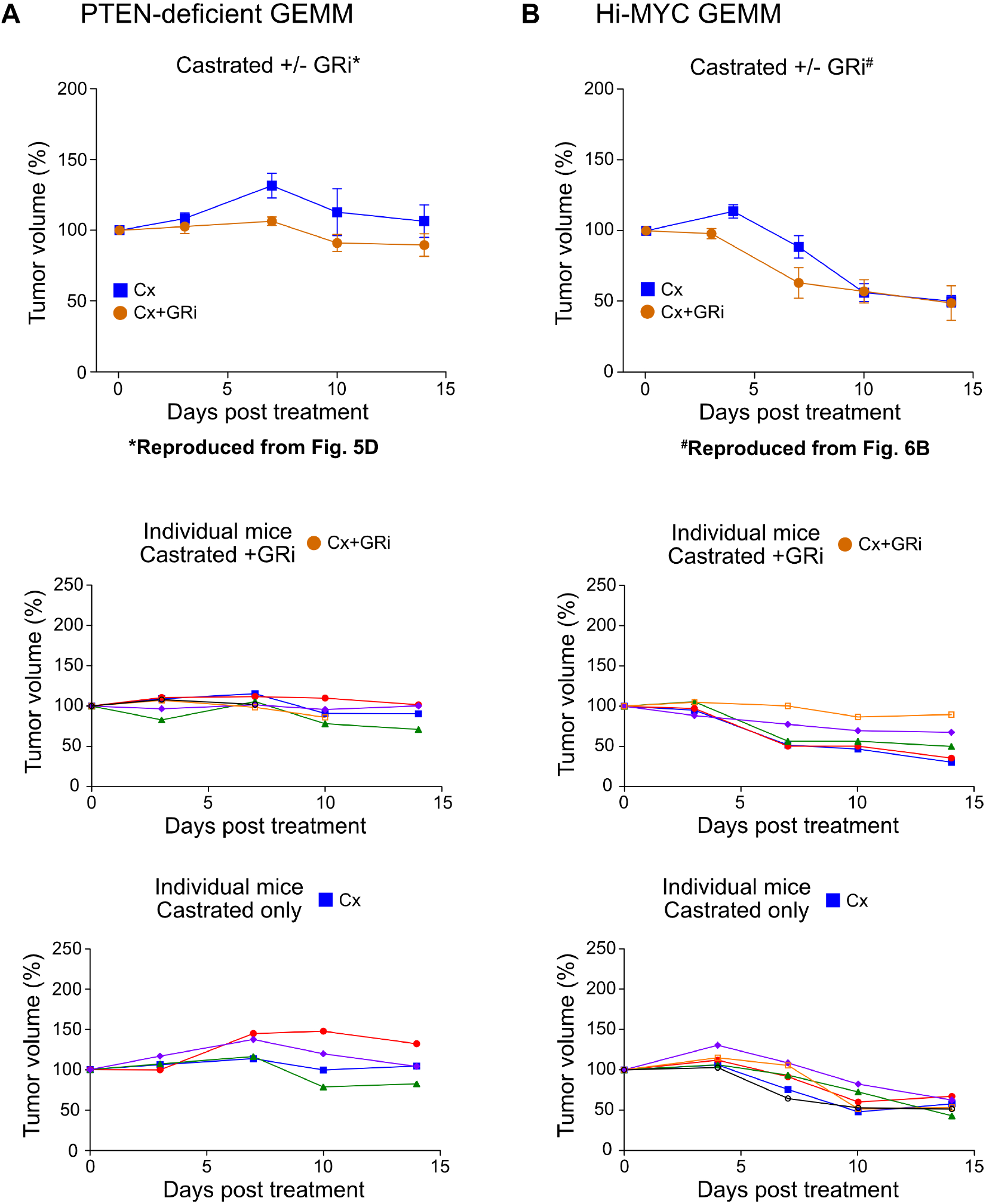
An inhibitor of glucocorticoid signaling blocks castration-induced delayed regression: plots of individual mice corresponding to Figures 5B and 6B. **A**. Relative tumor volume of PB-Cre4:Pten^*fl/fl*^ mice castrated and treated with/without CORT125281, as per Figure 5. Uppermost panel is a reproduction of Figure 5D (median +/-SE) ; the middle panel shows plots for individual castrated mice treated with CORT125281; and the bottom panel shows plots for individual castrated mice. **B**. Relative tumor volume of ARR_2_/PB-*Myc* mice castrated and treated with/without CORT125281, as per Figure 6. Uppermost panel is a reproduction of Figure 6B (median, +/- SE) ; the middle panel shows plots for individual castrated mice treated with CORT125281; and the bottom panel shows plots for individual castrated mice.

**Figure S4.**
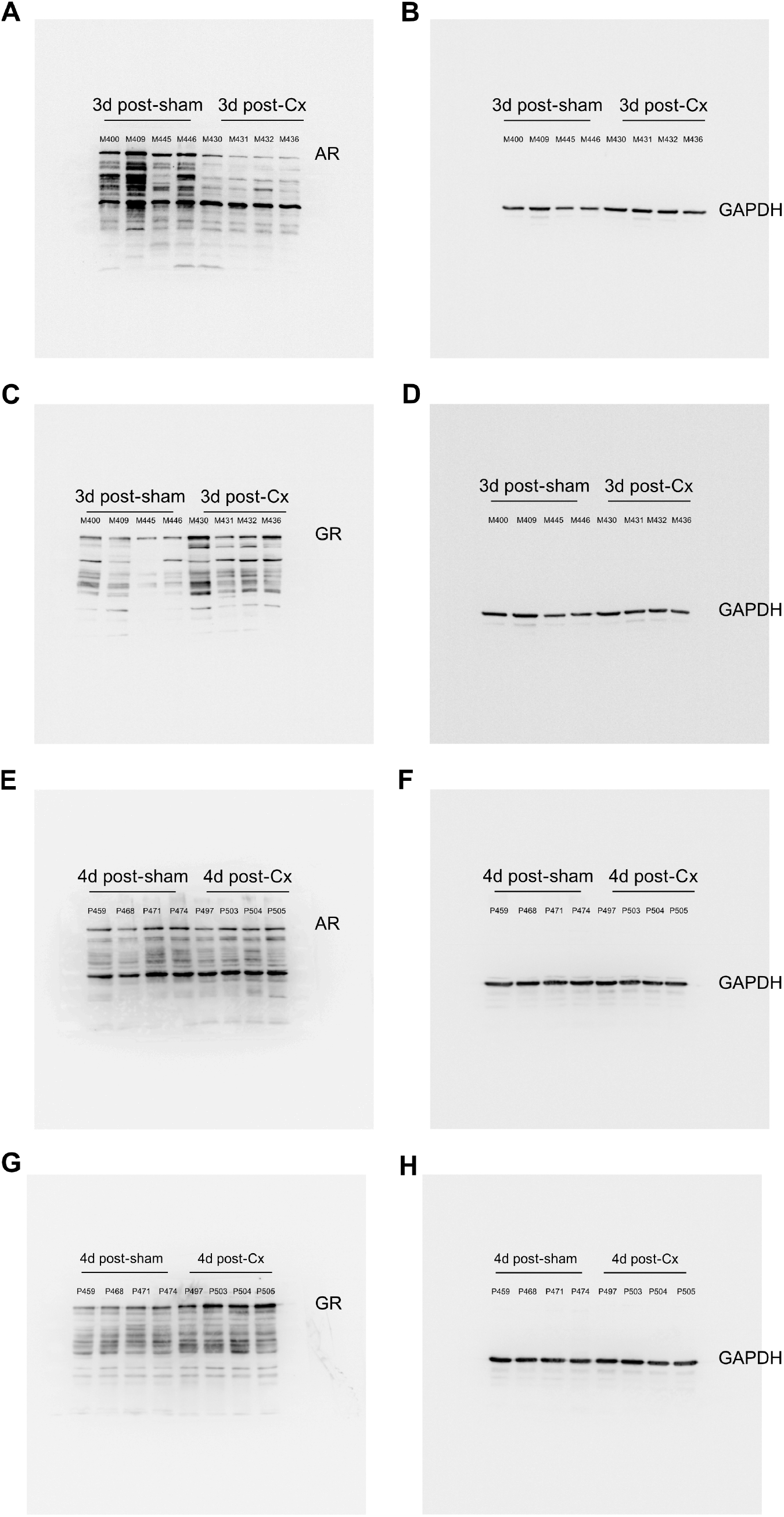
Complete immunoblot images corresponding to Figure 7. **A-D**. Immunoblots prepared using lysates from Hi- MYC GEMM (corresponding to Fig. 7B). **E-F**. Immunoblots prepared using lysates from PTEN-deficient GEMM (corresponding to Fig. 7A). Based on multiple publications and the mass markers, the full-length GR and AR proteins were identified as the slowest migrating band visible on the immunoblot image, as indicated.

**Figure S5.**
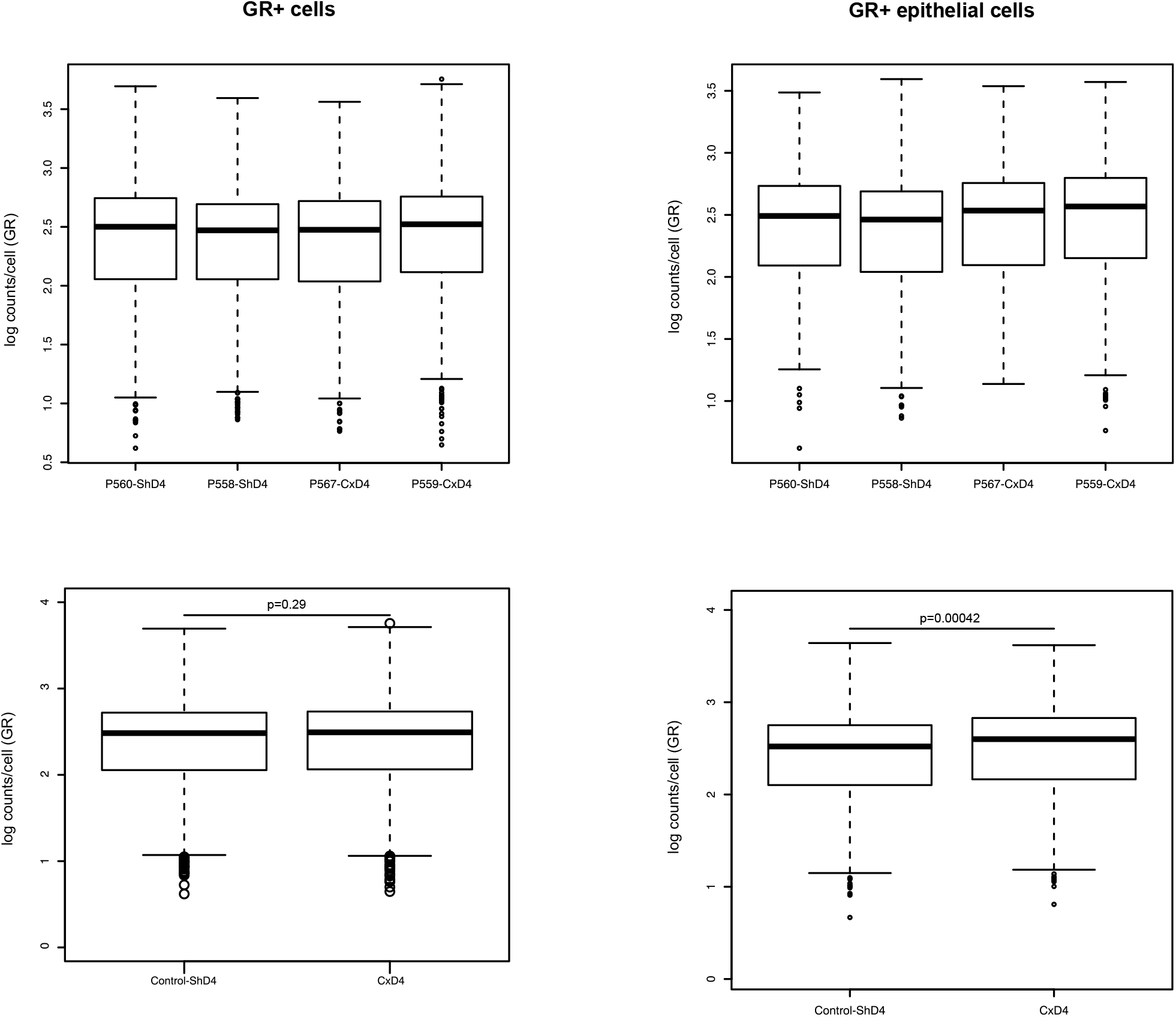
GR-transcript counts from scRNAseq data sets, corresponding to Figure 8C. Box plots of transcript counts corresponding to all GR-expressing cells (GR+ cells; left side) and in GR-expressing epithelial tumor cells (GR+ epithelial cells; right side), as identified in Figure 8A-B.The lower panels show the same data, with pooling of identically treated sets of mice. T-tests were performed on the pooled data and the p-values indicated in the figure.

**Figure S6.**
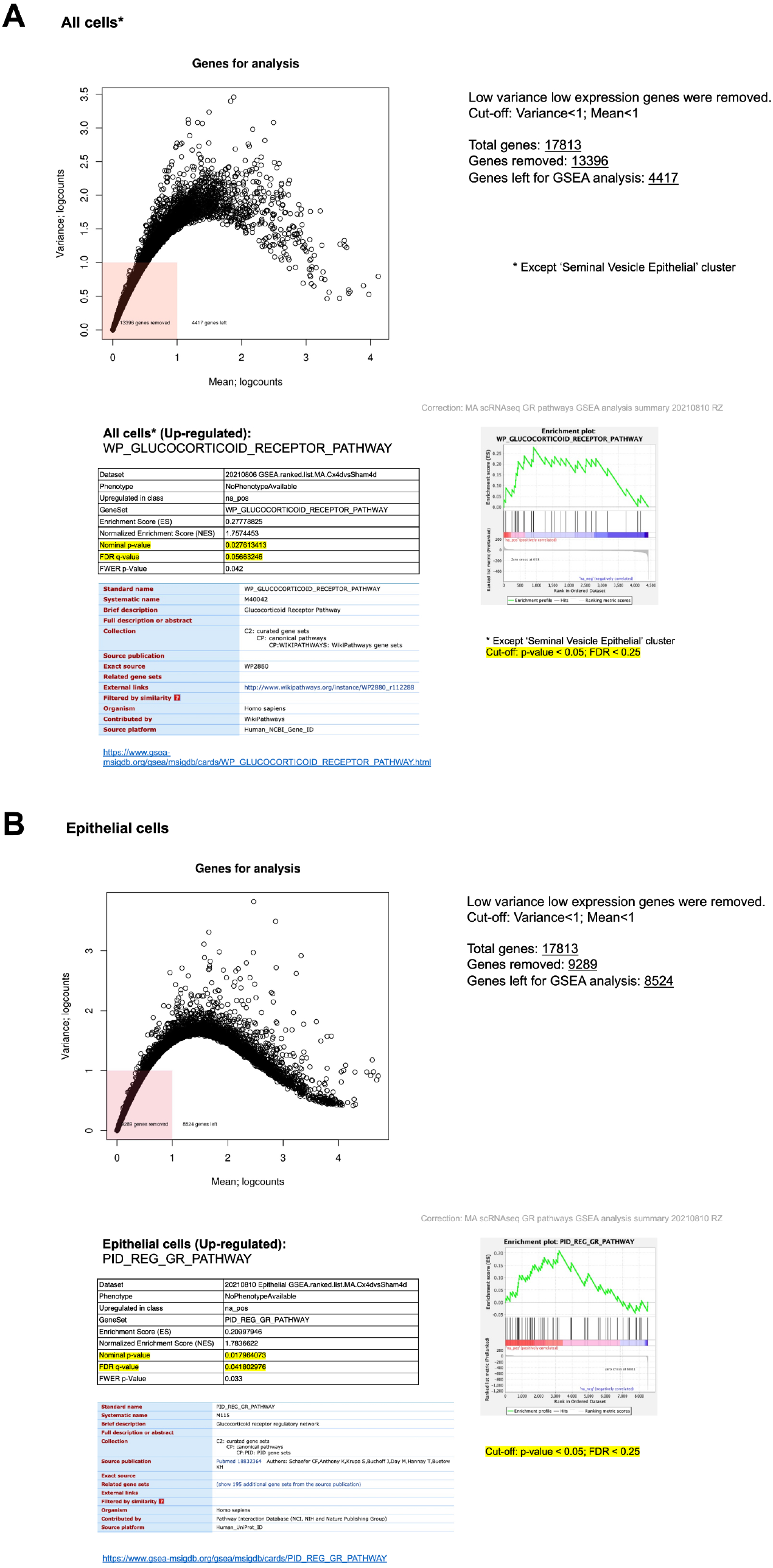
Extended data from differential gene expression and gene set enrichment analysis, corresponding to Figure 8D. Differential gene expression of the pooled transcripts from ‘All cells’ (excluding the seminal vesicle population) (A) and from the epithelial cell population only (B), followed by GSEA identified one glucocorticoid signaling gene set (PID_REG_GR _PATHWAY) that was significantly upregulated in the epithelial cell population. Genes with low variance and low expression across samples were removed before downstream analysis. **A**. All cells. Out of 17813 genes, 4417 genes (24.8%) were used to perform DEG across the entire cell population shown in Fig. 8A (excluding seminal vesicle). Among GR-related gene sets, only WP_GLUCOCORTICOID _RECEPTOR_PATHWAY was significantly upregulated in the 4d castrated group. **B**. Epithelial (tumor) cells only. Out of 17813 genes, we kept 8524 genes (47.9%) were used to perform DEG across the epithelial cell population shown in Fig. 8A. PID_REG_GR_ PATHWAY was significantly upregulated in castrated group. The enrichment plot is also shown in Fig. 8D

